# Naked mole rats have distinctive cardiometabolic and genetic adaptations to their underground low-oxygen lifestyles

**DOI:** 10.1101/2023.05.19.541451

**Authors:** Chris G. Faulkes, Thomas R. Eykyn, Jan Lj. Miljkovic, Rebecca L. Charles, Hiran A. Prag, Nikayla Patel, Daniel W. Hart, Michael P. Murphy, Nigel C. Bennett, Dunja Aksentijevic

**Affiliations:** School of Biological and Behavioural Sciences, Fogg Building, Mile End Road, Queen Mary University of London, London E1 4NS, UK; Department of Imaging Chemistry and Biology, School of Biomedical Engineering and Imaging Sciences, King’s College London, St Thomas’ Hospital, London, SE1 7EH, UK; MRC Mitochondrial Biology Unit, University of Cambridge, Keith Peters Building, Cambridge, CB2 0XY, UK; Centre for Clinical Pharmacology, William Harvey Research Institute, Bart’s and the London Faculty of Medicine and Dentistry, Queen Mary University of London, London, EC1M 6BQ, UK; Department of Medicine, University of Cambridge, Cambridge Biomedical Campus, Cambridge, CB2 0QQ; Centre for Biochemical Pharmacology, William Harvey Research Institute, Bart’s and the London Faculty of Medicine and Dentistry, Queen Mary University of London, London, EC1M 6BQ, UK; Department of Zoology and Entomology, Mammal Research Institute, University of Pretoria, Pretoria, 0002, South Africa

**Author notes:** **Corresponding author:** Dr Dunja Aksentijevic, Centre for Biochemical Pharmacology, William Harvey Research Institute, Barts and the London Faculty of Medicine and Dentistry, Queen Mary University of London, Charterhouse Square, London, United Kingdom, E. authors contributed equally (Chris G. Faulkes, Thomas R. Eykyn, Jan Lj. Miljkovic).

## Abstract

The naked mole-rat *Heterocephalus glaber* is a eusocial mammal exhibiting extreme longevity (37-year lifespan), extraordinary resistance to hypoxia and absence of cardiovascular disease. To identify the mechanisms behind these exceptional traits, RNAseq and metabolomics of cardiac tissue from naked mole-rats was compared to other African mole-rat genera. We identified metabolic and genetic adaptations unique to naked mole-rats including elevated glycogen, thus enabling glycolytic ATP generation during cardiac ischemia. Elevated normoxic expression of HIF-1α was observed while downstream hypoxia responsive-genes were down regulated, suggesting adaptation to low oxygen environments. Naked mole-rat hearts showed reduced succinate build-up during ischemia and negligible tissue damage following ischemia-reperfusion injury. These adaptive evolutionary traits reflect a unique hypoxic and eusocial lifestyle that collectively may contribute to their longevity and health span.

**One Sentence Summary:** Naked mole-rats have metabolic adaptations distinct from other subterranean genera rendering them resistant to cardiovascular pathology.

## Introduction

Naked mole-rats (*Heterocephalus glaber -*NMRs) are characterised by their social insect-like (eusocial) behaviour and a suite of adaptations to living in a hypoxic (low oxygen, O_2_) subterranean environment (*1*). The NMR (Fig. 1A) is one species within a clade of more than 30 species of African mole-rats, all of which adopt a subterranean lifestyle across sub-Saharan Africa within differing soil types and degrees of sociality (from solitary to eusocial, Fig. 1B,C) (*2-7*). Cooperative breeding and eusociality has evolved convergently within the African mole-rat clade. However, the NMR’s ancestral lineage diverged from their common ancestor around 30 million years ago (*8*). Despite similarities in their phylogeny and subterranean lifestyle, it is accepted that NMRs have many unique aspects to their biology compared to other African mole-rat species (*1*).

**Figure 1.**
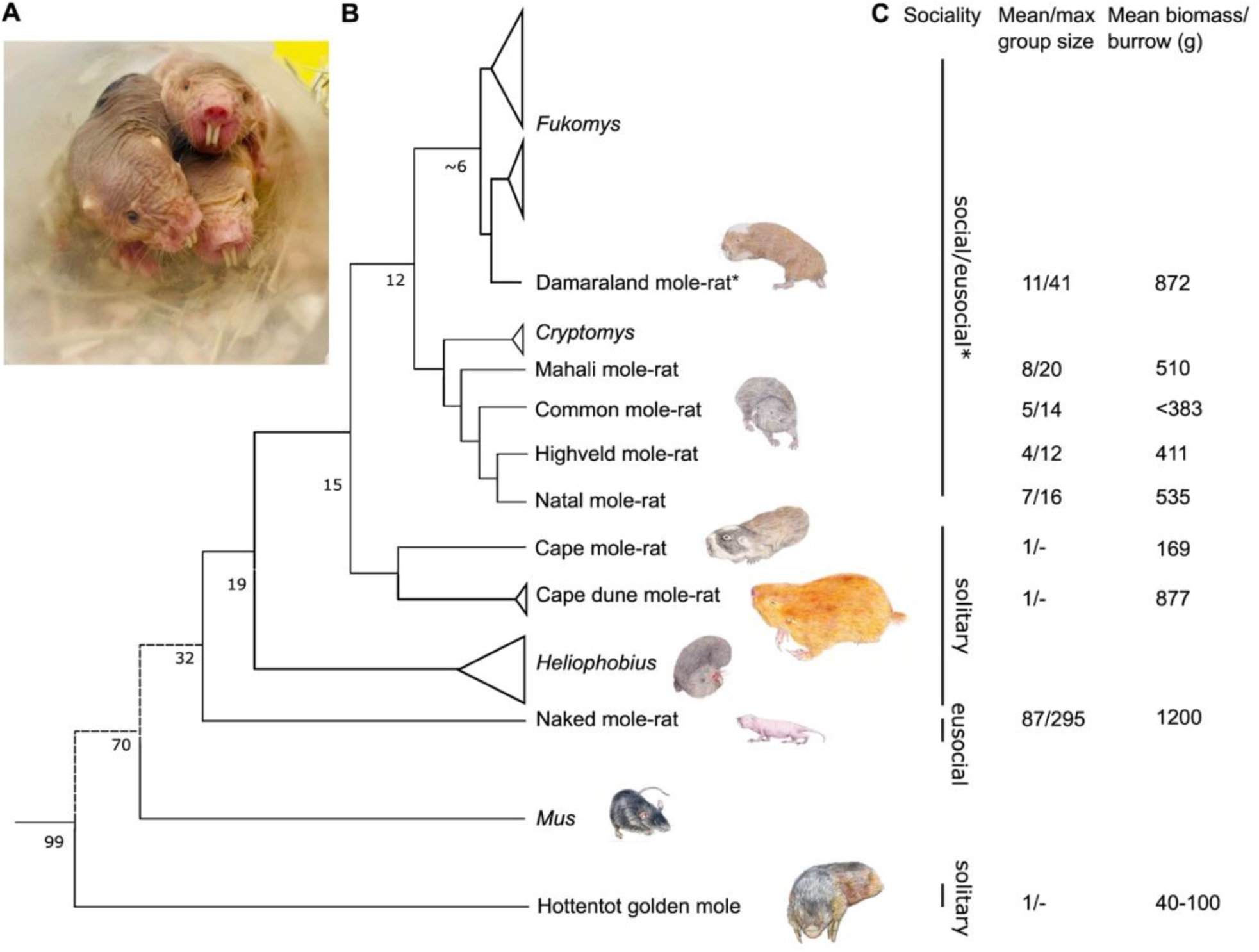
phylogenetic relationships and social status of African mole-rats, the mouse and the hottentot golden mole. (**A**) Naked mole-rats (*Heterocephalus glaber*) from the captive colonies at Queen Mary University of London. (**B**) Simplified molecular phylogeny for the African molerats indicating main clades/genera and species sampled in this study, together with the mouse *Mus* and the hottentot golden mole, *Ambysomus hottentotus*. Mole-rat phylogenetic relationships are based on mitochondrial 12S rRNA and cytochrome b sequence data, and analysis of 3999 nuclear genes. Numbers on internal nodes represent approximate divergence times in millions of years ago (myr). Adapted from (*11*). (**C**) Social lifestyles, mean and maximum social groups sizes and mean biomass of animals per burrow as indicated (*2-7*).

Up to 300 NMRs live together in an extensive burrow composed of labyrinths of underground tunnels, toilet chambers and a communal nest dug in fine, hard-packed soil, foraging for their staple diet of roots and tubers. Characteristically, mole-rats huddle together in nest chambers that are often many metres distant from transient mole-hills which open to the surface and can be up to 0.9 m deep (*9*). In this environment oxygen (O_2_) regularly becomes scarce and carbon dioxide (CO_2_) levels become elevated (*10, 11*). Out of all the species of African mole-rats, the potential for hypoxia as well as CO_2_ accumulation is greatest in NMRs as they have the longest burrows (3-4 km total tunnel length), the largest absolute group sizes and colony biomass that exceeds the other mole-rat species (Figure 1C) (*2-7*). Collectively, these living conditions limit NMRs gaseous exchange with the surface world. While data on O_2_/CO_2_ levels in wild NMR burrows is limited (*10*) and has never been measured in a nest chamber full of animals, NMRs in captive colonies are able to tolerate hours of extreme hypoxia (5% for 300 minutes) and can even survive up to eighteen minutes of complete oxygen deprivation (*12*). NMRs often elect to spend more time in areas of the burrow system with extreme atmospheric conditions including the nest chamber, where they may spend up to 70% of their time (*13*). This is something not regularly seen in other social African mole-rat species.

This challenging hypoxic habitat creates strong pressures and has been conducive to the evolution of unique adaptive traits in NMRs. Mammalian cells are not usually hypoxia-resistant, requiring uninterrupted O_2_ availability for survival. Fluctuations in O_2_ availability can lead to ischemia/reperfusion type tissue injury and irreversible organ damage. This is observed following a heart attack (*14*). Given the absence of cardiovascular disease in NMRs, despite regular fluctuating exposure between hypoxia/anoxia and normoxia, NMR hearts appear to have evolved resistance to both reduced O_2_ availability and ischemia/reperfusion (I/R) injury. NMR metabolism is known for its unusual features such as the ability to switch from glucose to fructose-driven glycolysis in the brain during anoxia (*12*). However, the mechanisms that underpin the extraordinary physiological adaptation to limited O_2_ availability in the heart are unknown. To determine how these adaptations arise in NMR, we hypothesised that the comparison to other African mole-rat genera would enable us to infer the changes in gene expression and metabolic signatures that contribute to the extreme hypoxia tolerance, resistance to cardiovascular injury and longevity of NMRs.

To do this, we performed RNAseq, metabolomics and pathway enrichment analysis on cardiac tissue from NMR (*Heterocephalus glaber*) and from seven other members of the African mole rat genera, Cape mole-rat (*Georychus capensis*), Cape dune mole-rat (*Bathyergus suillus*), Common mole-rat (*Cryptomys hottentotus hottentotus*), Natal mole-rat (*C. h. natalenesis*), Mahali mole rat (*C. h. mahali*), Highveld mole-rat (*C. h. pretoriae*) and Damaraland mole-rats (*Fukomys damarensis*) representing differing burrow and soil types, degrees of sociality, lifespan and hypoxia tolerance.

Morphological and life history characteristics of the African mole rat genera used in this study are summarised in Table S1. In addition, we include the evolutionarily highly divergent hottentot golden mole *(Ambysomus hottentotus*), an Afrotherian subterranean, solitary mammal. The African mole-rats and the Afrotherian clade last shared common ancestry around 100 million years ago (*8*) (Figure 1B). Therefore, the golden mole outgroup comparison should highlight similar adaptations to the subterranean niche. Moreover, the hottentot golden mole is sympatric with *Cryptomys* and can sometimes utilise their abandoned burrows, thus offering an outgroup that is subject to similar environmental conditions.

We show that unique cardiac metabolic and genetic features characterise the NMR heart compared to other African mole-rat genera and to evolutionarily divergent mammals, which result in tolerance to hypoxia/anoxia and lead to negligible tissue damage by I/R injury. These adaptations include supra-physiological glycogen content providing a readily available biochemical energy source via carbohydrate catabolism during anoxia/hypoxia, enhanced intracellular energy reserve (creatine) and constitutive stabilisation of HIF-1α under normoxic conditions. In addition, they show reduced succinate accumulation during ischaemia, which is a key metabolic source that drives the production of reactive oxygen species (ROS). NMRs express genetic adaptations at the mitochondrial level, which help to dampen ROS-related damage caused by fluctuating environmental O_2_ availability.

## Results

### NMRs have evolved a distinct cardiac gene expression profile

NMRs have evolved a cardiac gene expression profile that is *distinct* from any of the other mole rat genera as well as from the golden mole (Fig. 2A, Fig. 2B) and the C57/BL6 mouse (Fig. 2D), indicative of extensive adaptations. This is evidenced by separate clustering of NMR transcriptome data in the hierarchical cluster dendrogram of differential gene expression (Fig. 2A) and in the heatmap summary of predicted differentially expressed genes with the ability to encode transcription factors (Fig. 2B).

**Figure 2.**
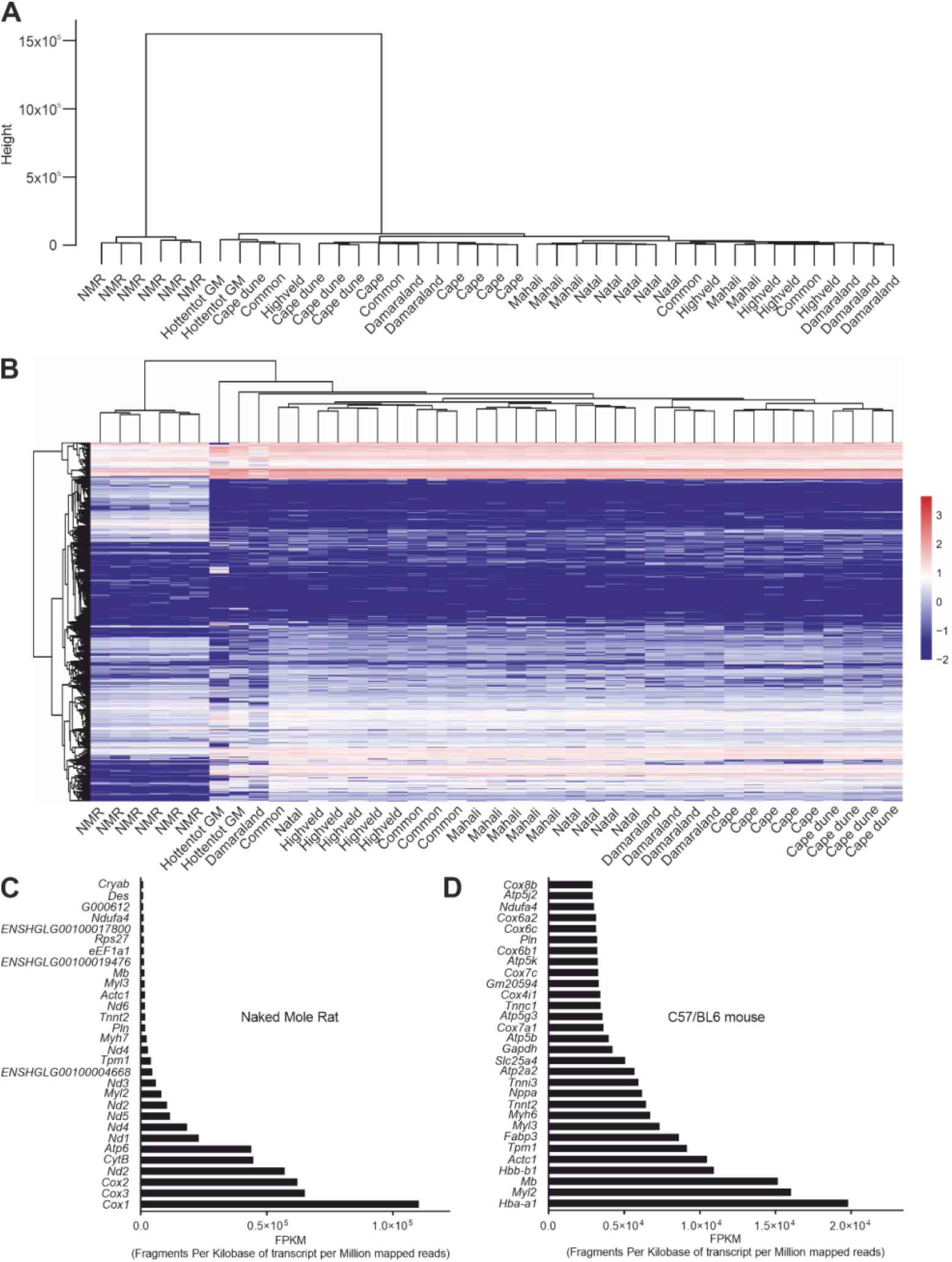
Naked mole rats exhibit distinct cardiac gene profile expression unrelated to any of the other subterranean genera and genetic adaptations at the mitochondrial level. (**A**) Hierarchical clustering of samples by similarity based on the expression level of all genes. (**B**) Heatmap with predicted differentially expressed genes encoding transcription factors (*TF)*. Normalised comparison of the 30 most expressed genes in (**C**) NMR hearts and (**D**) C57/BL6 mouse hearts – expressed as fragments per kilobase of transcript per million mapped reads. Nd2* and Nd4* in (C) are smaller novel transcripts (BGI_novel_G000346 and BGI_novel_G001276 respectively; Data S2)

Analysis of the 30 most abundantly expressed genes in NMR hearts vs C57/BL6 (Fig. 2C-D) highlights the importance of adaptations in energy metabolism. In NMR the majority, including the 10 most abundantly expressed, are mitochondrial oxidative phosphorylation complex genes (*Cox1, Cox3, Cox2, Nd2, Atp6, CytB, Nd1, Nd4, Nd3, Nd5, Nd6*, Fig. 2C). In contrast, in C57/BL6 mouse (*Mus musculus*) hearts the most abundantly expressed genes are responsible for myocardial contractile apparatus (i.e. myosin chains) and oxygen supply (i.e. haemoglobins) (e.g. *Hba a1, Myl2, Hbb-b1, Actc1*, Fig. 2D).

Comparison of the top 25 expressed cardiac genes across all the species analysed exemplifies the unique gene expression pattern of NMRs. Only 13 genes listed in the NMR top 30 are found in the top 30 of the other species, and the top 10 in the NMR are not shared across the other species (Data S1).

Pairwise comparisons of the RNAseq data from NMR compared to other mole rat genera are shown in the volcano plots in Fig. 3 (see Data S2 for a full list of differentially expressed genes in NMRs versus other mole-rats and the golden mole). Consistently higher expression of mitochondrial genes for the electron transport chain enzymes (OXPHOS) is observed in the NMR compared to the other subterranean species we examined. The following genes are all elevated: *Mettl12* (mitochondrial matrix methyltransferase protein 12)(*15*), *Cox1, Cox2* and *Cox3* (Cytochrome C oxidase, Complex IV); *Nad1*-*Nad6* (NADH dehydrogenase 2; Complex I); *CytB* (Cytochrome-b; Complex III) and *Atp6* (ATP synthase). Interestingly, Complex II (e.g., *Sdha-d*) was not differentially expressed in NMRs (Fig. 3, Fig. S3). Protein expression of complexes I, II, III and V was performed by western blot analysis (Fig. S3). Expression levels of complexes I, II and V were not found to be significantly different between NMR and C57/BL6 mouse while expression of complex III was elevated in NMR compared to C57/BL6 mouse.

**Figure 3.**
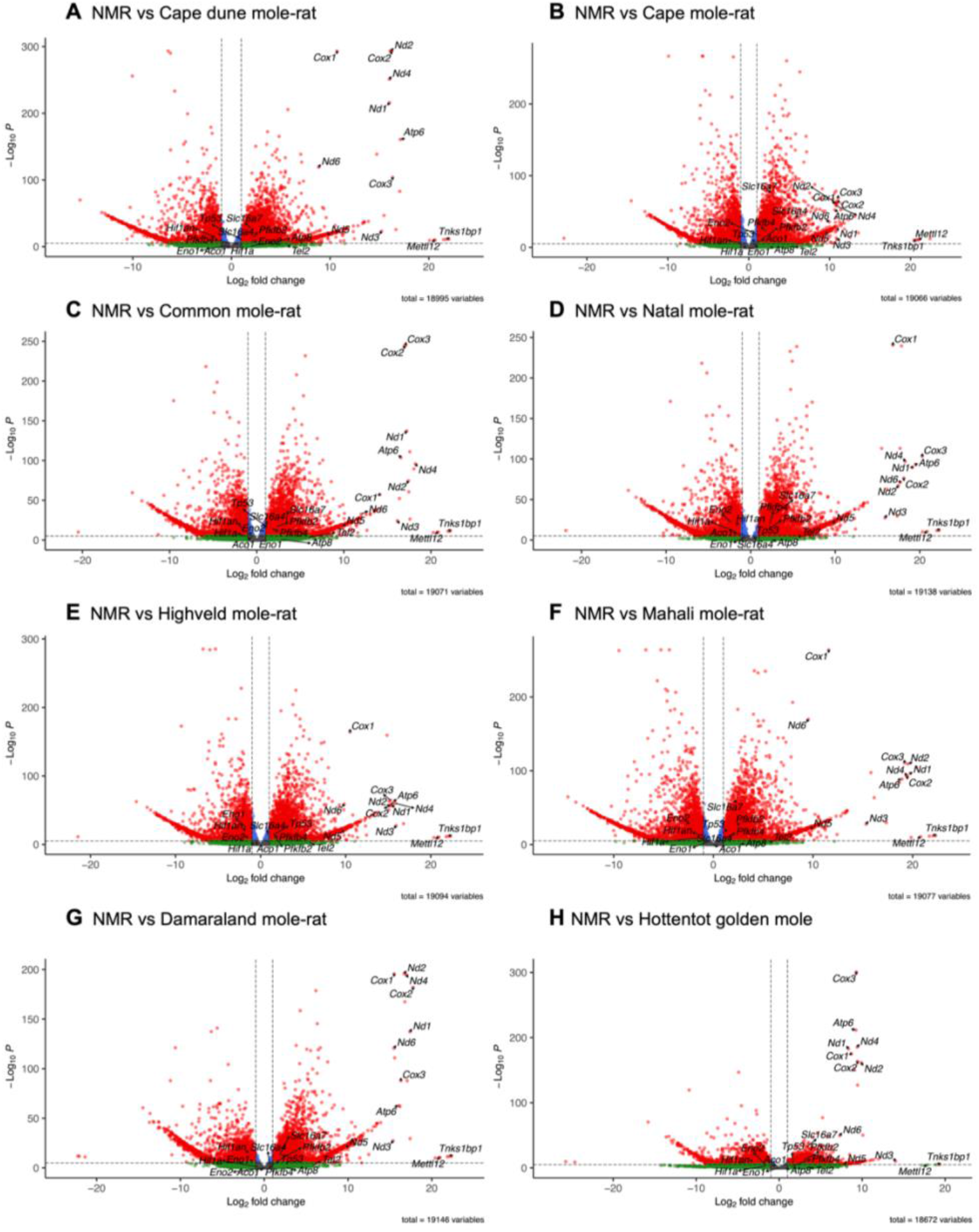
Volcano plot showing differentially expressed genes for each pairwise comparison between the naked mole-rat (NMR) and the other species in the RNA seq dataset (A-H). Y-axis shows statistical significance as the -Log_10_P values while the x-axis the Log_2_ fold change in gene expression (positive values represent overexpression in the NMR). Relevant genes that were identified and discussed in the text are labelled. The default cut-off for log_2_FC is > |2| and the default cut-off for P value is 10^−6^, denoted by the dotted lines. Points are coloured as follows: grey (not significant P > 10^−6^, log_2_FC < |2|), green (not significant P > 10^−6^, log_2_FC > |2|), blue (significant P < 10^−6^, log_2_FC < |2|), red (significant P < 10^−6^, log_2_FC > |2|).

### NMRs have evolved a distinct cardiometabolic profile

To better understand comparative genetic adaptations, semi-targeted metabolomic profiling was performed by high resolution ^1^H nuclear magnetic resonance spectroscopy on cardiac tissue from the NMR and from the other members of African mole rat genera as well as wild-type C57/BL6 mice (Fig. 4). NMRs showed a distinct cardiac metabolomic profile compared to other African mole-rat genera and the hottentot golden mole and mouse hearts, as shown in heatmap and hierarchical cluster summary (Fig. 4A). Principal Component Analysis (PCA) and Linear Discriminant Analysis (LDA) of cardiac metabolomic data showed clear separation between three groups (Fig. S2, Fig. 4B), the NMR being distinct, the other African mole rat genera clustering together suggesting they are metabolically far more similar to each other than to NMRs.

**Figure 4.**
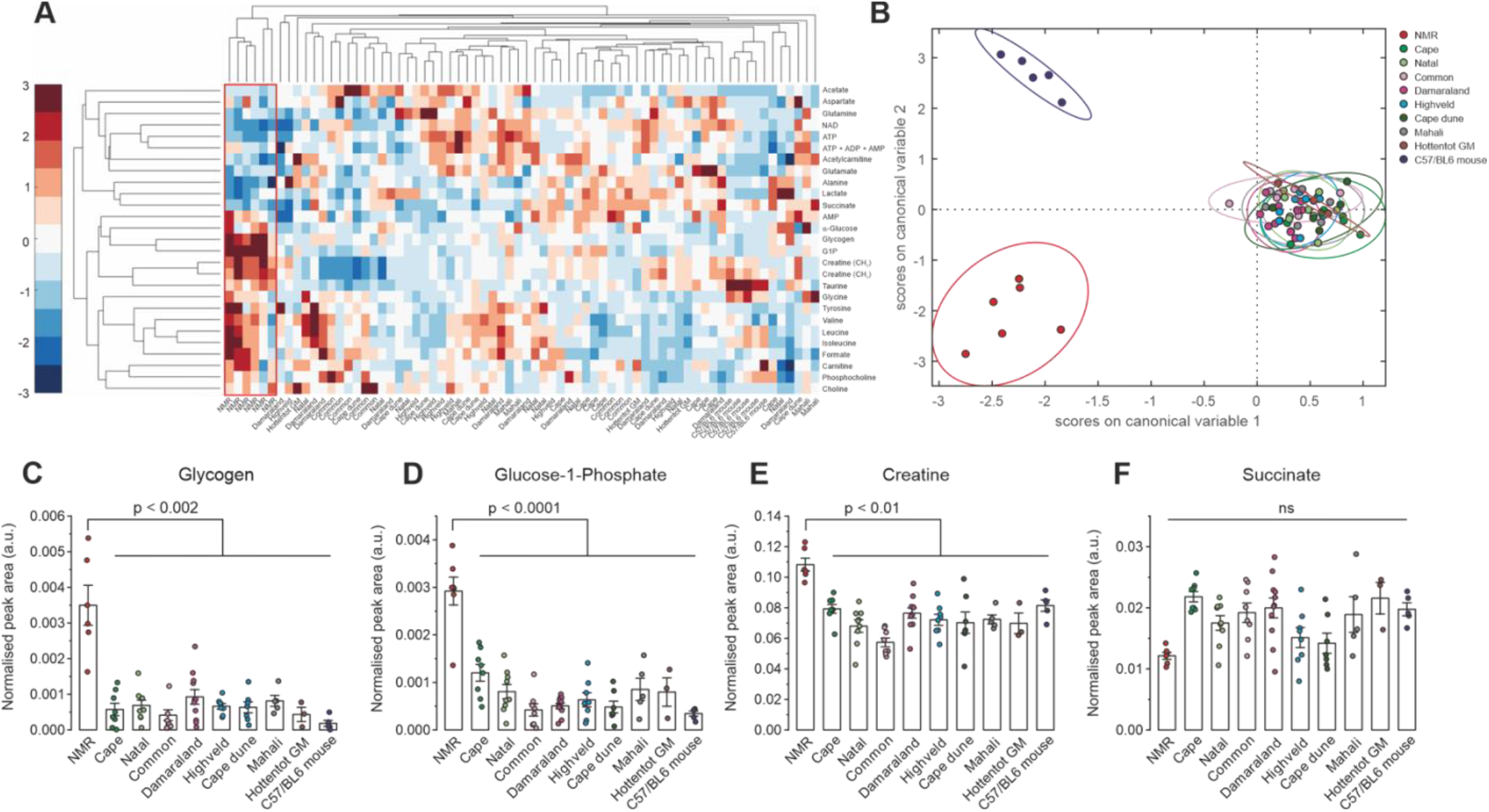
Enhanced cardiac energy reserve is unique to naked mole rat. **(A)** Heatmap summary of the cardiac metabolite concentrations determined by high resolution ^1^H nuclear magnetic resonance spectroscopy showing distinct profile of naked mole rat (NMR) versus cape, natal, common, damaraland, highveld, cape dune, mahali, the evolutionarily divergent hottentot golden mole or a mouse. Dendrograms top and left side show hierarchical cluster analysis of related genotypes (columns) and related metabolites (rows). **(B)** Linear discriminant analysis of the metabolite data in panel A showing clear separation of NMR from all other genera. **(C)** NMR is characterised by supra-normal myocardial glycogen content, (**D**) elevated glucose-1-phosphate, the product of glycogen catabolism, (**E**) elevated cellular energy reserve creatine, while succinate (**F**) was not significantly different under baseline normoxic conditions. t-test, data normality tested by Shapiro-Wilk. *P<0.05, **P<0.01 *** P<0.001 *** P<0.0001. (n=3-11/group).

The most notable metabolic difference observed between NMR and all other species was a significantly elevated myocardial concentration of glycogen (Fig. 4C), and of glucose-1-phosphate (Fig. 4D) which results from glycogen degradation; these were lower or undetectable in the other mole-rat species and undetectable in the mouse heart. Plasma metabolite analysis shows no evidence of increased insulin or circulating carbohydrates (Fig. S4). Thus, increased myocardial glycogen content in NMRs was not driven by altered exogenous metabolite and hormone availability and suggests a much greater capacity for storing myocardial glucose which can later be metabolised to generate energy. NMRs also had 44% elevation in myocardial creatine content compared to other mole-rat genera and C57/BL6 mouse (p<0.01, Fig. 4E). This is a remarkable increase in energy reserve in a healthy, non-transgenically modified heart.

### NMRs display constitutive stabilisation of HIF-1α under normoxic conditions

Another unexpected finding from our study is the evidence of stabilised cardiac HIF-1α protein expression in NMRs during normoxia (Fig. 5A-C, representative blots Fig. S5). Quantification of HIF-1α protein expression was carried out using three different antibodies against three different protein epitopes (Fig. 5) and was found to be significantly increased in NMR vs C57/BL6 mouse hearts (52% increase, average across three antibodies, Fig 5A-C, P<0.001). Stabilisation of HIF-1α expression under normoxia is surprising and contrasts with other organisms and tissues under normal O_2_ availability. Protein expression of lactate dehydrogenase (LDH) was elevated in NMR vs C57/BL6 mouse hearts, consistent with it being a target for HIF-1α (Fig. S3). Monocarboxylate transporter-1 (MCT-1) was also overexpressed in NMR compared to C57/BL6 mouse hearts (Fig. S3).

**Figure 5.**
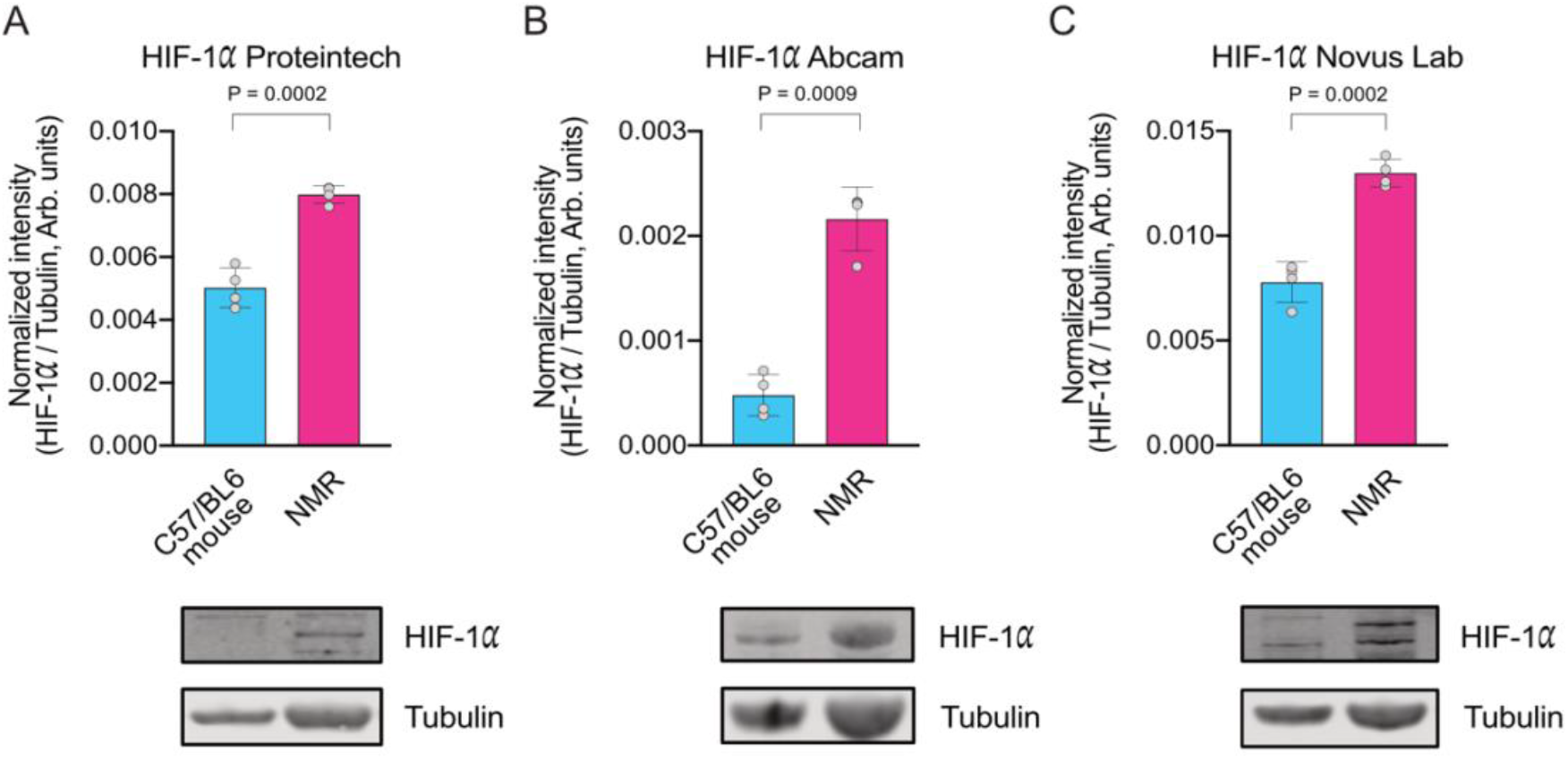
NMRs display constitutive stabilisation of HIF-1α under normoxic conditions. Quantification of HIF-1α protein expression using different antibodies against three different protein epitopes (Representative western blots in Supplementary Figure 5). Experimental groups contain n = 4 animals and western blot analysis was performed using several technical replicates. HIf-1α protein expression normalised to tubulin. Statistical analysis was performed using unpaired t-test (with Welch’s correction). (**A**) Proteintech rabbit polyclonal anti HIF-1α antibody (Clone developed against protein sequence including amino acids 574 - 799 of the human HIF-1 alpha protein; GenBank: BC012527.2). (**B**) Abcam rabbit polyclonal anti HIF-1α antibody (Clone developed against recombinant fragment. This information is proprietary to Abcam and/or its suppliers). (**C**) Novus Lab rabbit polyclonal anti HIF-1α antibody (Clone developed against a fusion protein including amino acids 530 - 825 of the mouse HIF-1 alpha protein; Uniprot #Q61221)

### Metabolic and genetic adaptations in NMR result in resistance to ischaemia reperfusion injury

In order to examine the physiological significance and functional translation of the genetic and metabolic differences in NMR (Fig. 6A), isolated hearts were subjected to 20 min ischemia followed by 2 hours of reperfusion (Fig. 6B). At baseline, NMRs maintained cardiac function low as evidenced from significantly lower left ventricular developed pressure (LVDP 30.2 ± 2.4 vs. 73.4 ± 2.7 mmHg, P<0.0001, Fig. S6) and heart rate (HR 134 ± 23 vs. 237 ± 33 beats/minute P<0.05, Fig. S6) compared to C57/BL6 mouse hearts. This is consistent with *in vivo* echo and MRI functional assessment (*16*).

**Figure 6.**
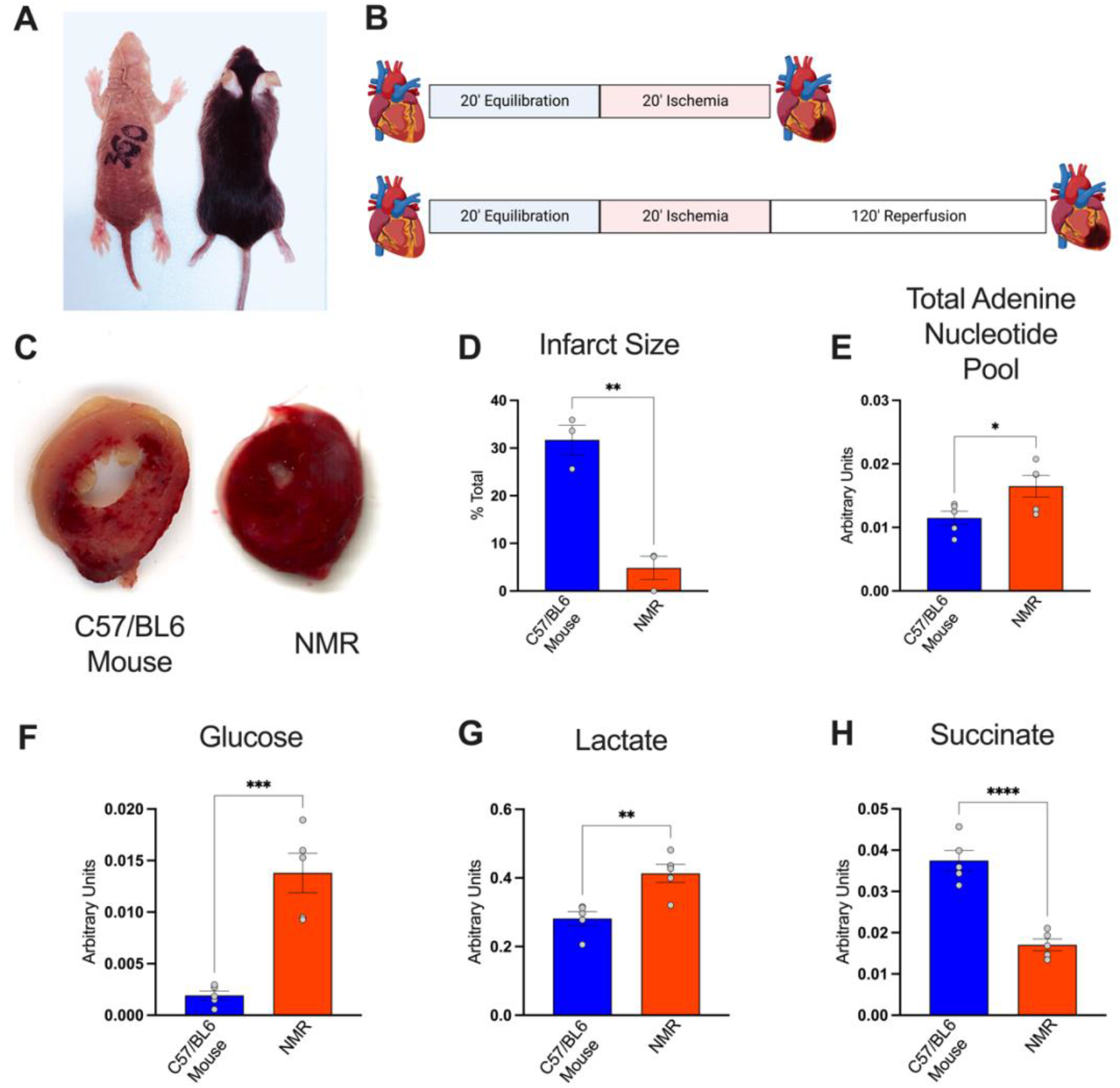
Naked mole-rat heart is resistant to ischemic damage. (**A**) Representative naked mole-rat and C57/BL6 mouse used in the study. (**B**) Langendorff heart ischemia/reperfusion protocols. (**C**) Representative cardiac sections images from C57/BL6 mouse and NMR post I/R. (**D**) Reduced infarct size in NMR vs C57BL6 post-myocardial ischemia (20 min) and reperfusion (120 min) protocol n=5/group, t-test, data normality tested by Shapiro-Wilk. *P<0.05, **P<0.01 *** P<0.001 *** P<0.0001. ^1^H nuclear magnetic resonance spectroscopy analysis of cardiac metabolites post-20-minute ischemia in NMR vs C57/BL6 mouse, (**E**) elevated total adenine nucleotide pool (sum of ATP+ADP+AMP), (**F**) elevated glucose, (**G**) elevated lactate and (**H**) reduced myocardial succinate.

After 20 minutes total global normothermic ischemic anoxia, NMR hearts had enhanced energy supply (29% higher total adenine nucleotide pool, 62% glucose, 52% lactate, Fig 6 E, F, G) and reduced succinate accumulation (54% lower, Fig. 6H), the metabolic source of ROS in I/R damage. NMR hearts showed minimal damage post 20-minute global normothermic ischemia as infarcted scar tissue was barely detectable unlike in mouse hearts which exhibited extensive tissue damage (Fig 6. D-G). Biochemical analysis of the tissue also revealed that the NMR hearts accumulate lactate during ischemia that effluxes immediately upon reperfusion (1-minute reperfusion, Fig. S7) and then returns rapidly to oxidative metabolism as the lactate levels 1-hour post-reperfusion are comparable to equilibration/pre-ischemia, as well as comparable to mouse heart (Fig. S7).

## Discussion

RNAseq and metabolomics were performed on cardiac tissue from NMR and from seven other African mole rat genera, the evolutionarily divergent hottentot golden mole and the C57/BL6 mouse. A plethora of metabolic and genetic characteristics were unique to the NMR. Principal component and hierarchical cluster analysis of RNAseq data uniquely separated NMR data from all other groups. There is a significant upregulation of mitochondrial genes for much of the electron transport chain enzymes including Complex I, III, IV as well as ATP synthase. Interestingly, succinate dehydrogenase was unaltered across genera. Additionally, Mct2, Pfkfb2, several glycolytic genes and the master tumour suppressor gene Tp53 were also consistently upregulated (Fig 3).

Taken together, the genetic adaptations are unique to NMR and not shared with other members of the African mole rat genera, thus, reflecting adaptive traits within the heart genome unique to the lifestyle of NMRs.

Principal component and discriminant analysis of metabolomic data also uniquely clustered NMR from all other groups. Compared to the other genera, metabolic adaptations included supra-normal glycogen content and G-1-P arising from enhanced glycogen turnover, thus providing readily available biochemical energy (ATP) via carbohydrate catabolism during stress. The concentration of myocardial glycogen is higher than other mammals and was higher than measured in the post-prandial liver where its concentration of this metabolite is the highest (NMR heart glycogen 1.2 ± 0.02 versus post-prandial mouse liver glycogen 0.4 ± 0.08 μmol g−1; p < 0.001) (*17*).

NMR also showed enhanced energy reserves in the form of elevated creatine. It has previously been shown that elevated creatine and glycogen are protective during myocardial ischaemia as they improve energy reserve (*19*). Increased myocardial creatine in creatine transporter overexpressing mice has been shown to be beneficial as it protects against I/R injury by decreasing necrosis in a dose-dependent manner (*19*). Transgenic hearts with elevated creatine exhibited improved functional recovery following *ex vivo* I/R (59% of baseline vs. 29% of baseline cardiac function) (*19*). Furthermore, cellular creatine loading has been shown to delay mitochondrial permeability transition pore opening in response to oxidative stress, suggesting an additional mechanism to prevent I/R injury (*19*). However, in other mammals it is not possible to increase myocardial creatine concentrations to supra-normal levels because it is subject to tight regulation by the sarcolemmal creatine transporter (*18*). Furthermore, artificial increases in myocardial creatine content by transgenesis have also shown that the hearts of other mammals are incapable of maintaining the augmented creatine pool adequately phosphorylated, resulting in increased free ADP levels, LV hypertrophy, and significant dysfunction (*18*). Taken together metabolic adaptations are unique to NMR and not shared with the other members of African mole rat genera, thus, these adaptations also do not construe mammal-wide convergence in adaptive traits within the heart metabolome.

NMR hearts showed a significant elevation of HIF-1α levels under normoxic conditions, compared to the C57/BL6 mouse. This was confirmed at the protein level using several different antibodies raised against different HIF-1α epitopes to best compare two different animal species. This phenomenon contrasts with what is currently known about hypoxia/anoxia-mediated degradation of HIF-1α protein. HIF-1α is an oxygen-dependent transcriptional activator that mediates a wide range of adaptations to reduced oxygen. Under normoxia HIF-1α expression in NMR is constituent (*20*) and stable due to the mutation in HIF-1α (T407 to I) and in Von Hippel–Lindau (VHL; V166 to I) that both synergistically prevent the ubiquitin-mediated degradation of HIF-1α (*20*).

Unlike humans who are prone to heart injury by hypoxia and anoxia caused by blood occlusion during heart attacks, NMR hearts appear to have adapted to evade such damage. Ischaemia or hypoxia typically leads to elevated levels of succinate in the heart, and other tissues, and is a key driver for the production of ROS during reperfusion (*14*). In the hearts of NMR subjected to ischemia, we observed lower levels of succinate than the mouse hearts. Low succinate levels may limit ROS production during reperfusion and therefore mitigate tissue damage.

However, succinate is an inhibitor of prolyl hydroxylase (PHD), and therefore reduced accumulation of succinate may also be insufficient to inhibit PHD, leading to weak stabilisation of HIF-1α and its enhanced degradation during ischaemia. Interestingly, blunted HIF-1α response in anoxia is also observed in human tissues that experience regular variation of O_2_ such as anoxic cord blood monocytes (*21*). In keeping with the reduced succinate, NMR hearts showed negligible tissue damage post ischaemia reperfusion, while the C57/BL6 mouse showed extensive tissue damage due to myocyte necrosis and reperfusion injury.

It has also been shown that the mitochondria from the whole host of tissues (skeletal muscles, diaphragm, heart, spleen, brain, lung kidney) from mice, bats and NMRs contain two mechanistically similar systems to prevent ROS generation: mitochondrial membrane bound hexokinases (I and II) and creatine kinase. Specifically, both of these metabolic systems function in a way that one of the kinase substrates (ie. mitochondrial ATP) is electrophoretically transported by the ATP/ADP antiporter to the catalytic site of either bound hexokinase or creatine kinase without affecting ATP concentration in the cytosol. ADP, another kinase reaction product, is transported back to the mitochondrial matrix *via* the antiporter by the same electrophoretic process. This system continuously supports ATP synthase independent of glucose and creatine availability. This unique ATP homeostatic mechanism would be supported by the elevated creatine and glycogen observed in NMR hearts at baseline (normoxic conditions) making them available for ATP provision during times of stress (ie. ischemia) (*22*). These conditions keep mitochondria in a state of mild depolarization with the membrane potential ΔΨ maintained at a lower than maximal level (*22*), sufficient to completely inhibit mROS generation. During aging in C57/BL6 mice (2.5years), mild depolarization disappears in most tissues including the heart, however, age-dependent decreases in the levels of bound kinases is not observed in NMRs (*22*). As a result, ROS-mediated protein damage, which is substantial during the aging of short-lived mice, is stabilised at low levels during the aging of long-lived NMRs. It has been suggested that mild mitochondrial depolarization is a crucial component of the anti-aging system (*22*) contributing to NMR longevity.

Our study provides insights into the evolution of the NMR’s remarkable cardiac adaptations to life in hostile conditions in the dark and at low O_2_. Collectively, they may contribute to their exceptional longevity and resistance to ischemic pathologies. The extreme metabolic and genetic traits we identified offer opportunities for advancing other areas of physiological and medical research, including the development of novel therapeutic approaches.

## Acknowledgements

We would like to thank Dr Harold Toms and Dr Nasima Kanwal QMUL NMR facility for the technical assistance with the NMR spectroscopy and Prof Yannick Wurm for constructive criticism of the study.

## Author contributions

Conceptualization: DA, CGF, Methodology: DA, CGF, TRE, Investigation: DA, CGF, TRE, JLM, RC, NP, HAP, DH, NB, Visualization: DA, CGF, TRE, JLM, Funding acquisition: DA, Project administration: DA, Supervision: DA, Writing – original draft: DA, CGF, TRE, JLM, Writing – review & editing: DA, CGF, TRE, JLM, MPM

## Data and materials availability

Data available on reasonable request from the corresponding author.

