## Supplemental Material for "Naked mole rats have distinctive cardiometabolic and genetic adaptations to their underground low-oxygen lifestyles"

**The PDF file includes:**

Materials and Methods
Supplementary Text
Figs. S1 to S7
Tables S1
References

### Materials and Methods

Animals

NMRs were bred at the School of Biological and Behavioural Sciences, Queen Mary University of London. The non-breeding adult NMRs used in this study were second-generation or more captive-born, descended from animals captured in Kenya in the 1980s. Colonies were maintained using artificial burrow systems (group-housed in interconnected multi-cage systems) at 30°C and 21% $O_2$ in 50% humidity with a 12h light cycle. Their diet consisted of fresh vegetables, fruit and tubers (sweet potatoes) *ad libitum*. The ages selected for this study allowed for physiological age matching such that all animals were at equivalent percentages of maximum lifespan and therefore not the same chronological age. C57/BL6 (J strain) mice were purchased from Charles River Laboratories, UK and kept in individually ventilated cages, with access to food and water *ad libitum*.

*Georychus capensis, Bathyergus suillus, Cryptomys hottentotus hottentotus, C. h. pretoriae, C. h. mahali*, and *C. h. natalenesis* were wild-captured in South Africa using Hickman live traps, baited with a small piece of sweet potato. All traps were monitored for captures every 2-3 hours over the course of the day and left overnight, being checked first thing in the morning. Permission to capture these species was obtained from all landowners, and a collecting permit was obtained from the relevant nature conservation authorities (Permit number: Western Cape- CN44-87-13780, Gauteng- CPF6-0124, Kwa-Zulu Natal- OP1545/2021). *Fukomys damarensis* were laboratory maintained. They were individually housed at the University of Pretoria at ~27°C and 21% $O_2$ in 50% humidity with a 12L:12D light cycle. The Animal Use and Care Committee of the University of Pretoria evaluated and approved the experimental protocol and collection of all samples (ethics clearance number: NAS022/2021), with DAFF section 20 approval (SDAH-Epi-21051907211). Animal experiments at QMUL were approved by the local ethical review committee and in accordance with the Home Office Animals in Scientific Procedures Act 1986.

Langendorff Heart Perfusion

Beating hearts were excised from terminally anaesthetized naked mole rats (n=5) and C57/BL6 mice (n=5) (Charles River, UK) for Langendorff perfusions as previously described (*1*). After 20 minutes of equilibration, hearts were subject to 20 minutes of global normothermic ischemia. Hearts were snap frozen at the end of the protocol using a Wollenberger clamp pre-cooled in liquid $N_2$. Ischemia/reperfusion (I/R) outcome in NMRs was compared to C57BL6 mouse heart due to their anatomical, physiological, and genetic similarity to humans. We were unable to repeat *ex vivo* cardiac perfusion protocols in all other 5 mole rat genera hearts as they are not kept in captive colonies in the UK.

### Myocardial infarct size quantification

Myocardial infarct size was quantified as described previously (*2*). In brief, after 20 min of equilibration, Langendorff perfused hearts (n=3/ group) were subject to 20 min of global normothermic ischemia and 2 hours of reperfusion. At the end of the protocol, hearts were perfused for 10 mins with 3% triphenyltetrazolium chloride (TTC) in KH Buffer followed by 10 min incubation in 3% TTC-KH. Tissue was sectioned (mouse heart gauge, Zivic instruments, USA) and infarct field was quantified using ImageJ Software (*2*).

### $^{1}$H NMR Spectroscopy

Powdered heart samples were subject to methanol / water / chloroform phase extraction (*1*). Frozen heart tissue was homogenised in 2 mL each of ice-cold methanol, chloroform, and Millipore water and vortexed. Samples were centrifuged for 1 hour at 3600 rpm at 4°C to separate aqueous, protein, and lipid layers. The upper aqueous phase was separated, 20-30 mg chelex-100 was added to chelate paramagnetic ions, vortexed and centrifuged at 3600 RPM for 1 minute at 4°C. The supernatant was transferred to a fresh falcon tube containing 10 µL of universal pH indicator solution followed by vortexing and lyophilization. Dual-phase-extracted metabolites were reconstituted in 600 µL of deuterium oxide [containing 8 g/L NaCl, 0.2 g/L KCl, 1.15 g/L $Na_2HPO_4$, 0.2 g/L $KH_2PO_4$ and 0.0075% w/v trimethylsilyl propanoic acid (TSP)] and adjusted to pH ≈ 6.5 by titrating with 100 mM hydrochloric acid.

$^{1}$H nuclear magnetic resonance spectra were acquired using a vertical-bore, ultra-shielded Bruker 14.1. tesla (600 MHz) spectrometer with a bbo probe at 298K using the Bruker noesygppr1d pulse sequence. Acquisition parameters were 128 scans, 4 dummy scans and 20.8 ppm sweep width, acquisition time of 2.6s, pre-scan delay of 4s, 90° flip angle and experiment duration of 14.4 minutes per sample (*3*). TopSpin (version 4.0.5) software was used for data acquisition and for metabolite quantification. FIDs were multiplied by a line broadening factor of 0.3 Hz and Fourier-transformed, phase and automatic baseline-correction were applied. Chemical shifts were normalised by setting the TSP signal to 0 ppm. Metabolite peaks of interest were initially integrated automatically using a pre-written integration region text file and then manually adjusted where required. Assignment of metabolites to their respective peaks was carried out based on previously obtained in-house data, confirmed by chemical shift and using Chenomx NMR Profiler Version 8.1 (Chenomx, Canada). Peak areas were normalized to the total metabolite peak area.

### Western blotting

Powdered heart samples were homogenised (100 µL of buffer per 10 mg of cardiac tissue) on ice in 100 mM Tris buffer (pH 7.4) supplemented with complete mini EDTA-free protease inhibitor (Roche) using a glass tissue grinder. The tissue homogenates were re-suspended in an equal volume of 2x reducing SDS sample buffer. Protein were resolved by SDS-PAGE (4 – 20 % Mini-PROTEAN TGX, Bio-Rad or Novex 4 – 20% Tris-Glycine, ThermoFisher Scientific) using Mini-Protean 3 system (Bio-Rad) or XCell4 SureLock Midi Cell and transferred using semi-dry system (Bio-Rad) to 22 µm PVDF or Nitrocellulose membranes (Bio-Rad). Following transfer, membranes were blocked for one hour with 5 % non-fat dried milk in TPBS at RT. After blocking, membranes were immunoprobed with primary antibodies dissolved in 5 % non-fat dried milk in TPBS overnight. Subsequently, membranes were washed three times with TPBS and incubated with IR-dye conjugated secondary antibodies in Intercept blocking buffer (LI-COR) or with horseradish peroxidase-coupled anti-rabbit IgG antibody in 5 % non-fat dried milk in TPBS for one hour at RT in dark.

After the incubation, membranes were washed three times with TPBS and imaged using Odyssey DLx infrared imaging system (LI-COR) or ECL reagent (ThermoFisher Scientific) and imaging cabinet. Protein bands were analysed and quantified using Image Studio Lite (LI-COR) or ImageJ (NIH).

List of vendors and dilution of antibodies used in this study:

Anti HIF-1α Proteintech (#20960-1-AP); (Clone developed against protein sequence including amino acids 574 - 799 of the human HIF-1α protein; GenBank: BC012527.2) 1: 1000
Anti HIF-1α Abcam (#ab 179483); (Clone developed against recombinant fragment. This information is proprietary to Abcam and/or its suppliers) 1: 1000
Anti HIF-1α Novus lab (#NB100-479); (Clone developed against a fusion protein including amino acids 530 - 825 of the mouse HIF-1α protein; Uniprot #Q61221) 1: 1000
Anti β-Tubulin ThermoFisher Scientific (#62204) 1: 5000
Anti α-Actin Merck Millipore (#MAB1501) 1: 3000
Anti Vinculin Sigma Aldrich (#SAB4200729) 1: 5000
Total OXPHOS Rodent WB Antibody Cocktail (#ab110413) 1: 1000
Anti LDH Abcam (#ab 74010) 1: 5000 (antibody cross-reacts with LDHA, B and C)
Anti MCT-1 Proteintech (#20139-AP) 1: 1000

RNAseq/transcriptome analysis

Total RNA extraction and analysis was performed commercially by BGI Tech solutions (Hong Kong) using sequencing platform DNBseq and read length PE 100bp. We sequenced 41 samples as follows: Naked mole-rat, *Heterocephalusglaber* (n=6), Cape mole-rat, *Georychus capensis* (n=5), Cape dune mole-rat, *Bathyergus suillus* (n=4), Common mole-rat, *Cryptomys hottentotus hottentotus*, *C. h. natalenesis* (n=5), Mahali mole-rat, *C. h. mahali* (n=5), Highveld mole-rat *C. h. pretoriae* (n=5), Damaraland mole-rat, *Fukomys damarensis*, Hottentot golden mole, *Ambysomus hottentotus* (n=2) and C57BL6 mice *Mus musculus* (n=6). An average of 4.28 Gb of sequence was generated per sample.

Standard bioinformatic processing was performed as a service by BGI Tech Solutions. Briefly, the reads mapped to rRNAs were removed to produce rawdata, then low quality reads were filtered where more than 20% of the bases qualities were lower than 10, reads with adaptors and reads with unknown bases (N bases more than 5%) were also removed to produce "clean reads". Clean reads were mapped onto reference genome (*Heterocephalus glaber* Ensemble_release-108). Genome mapping was conducted using HISAT2 (Hierarchical Indexing for Spliced Alignment of Transcripts; http://www.ccb.jhu.edu/software/hisat).

The average mapping ratio with the reference genome was 18.13% while the the average mapping ratio for genes was 32.28%; 20,285 genes were identified. Due to limited mapping rate using the naked mole-rat as a reference for the comparison with the mouse C57BL6 transcriptome (and *vice versa*), it was not possible to do the same range of analyses for the mouse versus the naked mole-rat, as for the naked mole-rat versus the subterranean species listed above. Mapping was followed by novel gene prediction, SNP and INDEL calling and gene splicing detection. StringTie (http://ccb.jhu.edu/software/stringtie) was used to reconstruct transcripts, while Cuffcompare and Cufflinks tools (http://cole-trapnell-lab.github.io/cufflinks) were used to compare reconstructed transcripts to the reference annotation.

Novel transcripts were defined as (i) unknown, intergenic transcripts, (ii) transfrags falling entirely within a reference intron, (iii) generic exonic overlaps with a reference transcript and (iv) potentially novel isoforms (fragments) where at least one splice junction is shared with a reference transcript. CPC (http://cpc.cbi.pku.edu.cn) was then used to predict the coding potential of novel transcripts. Novel coding transcripts where then merged with reference transcripts to get a complete reference, and downstream analysis was based on this complete reference. GATK(*4*) was deployed to call SNPs and INDELs for each sample.

Finally, DEGs (differentially expressed genes) were identifed between samples with DEseq2 (*5*) and clustering analysis and functional annotations performed. Hierarchical clustering for DEGs was performed using pheatmap, a function of R. Following Gene Ontology Analysis of DEGs GO functional enrichment analysis was undertaken using phyper, a function of R. The analysis pipeline is summarised in Figure S1 below.

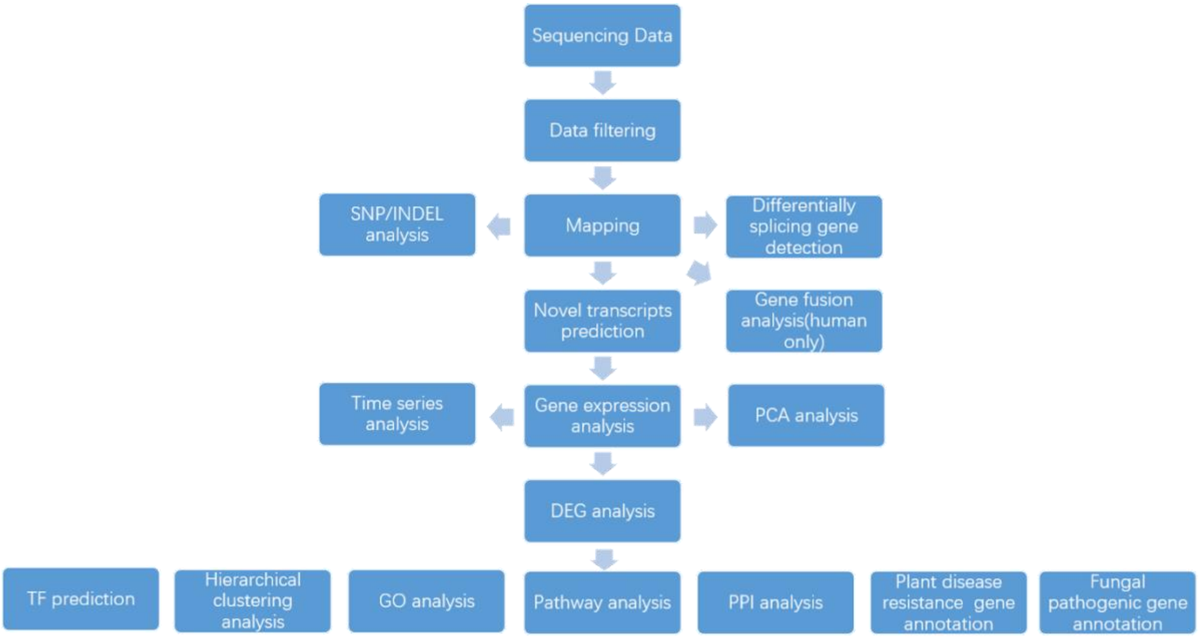


**Fig S1.** Summary of the bioinformatics workflow.

### Statistical Analysis

Data are presented as mean ± SEM. Comparison between groups was performed by Student's t-test (Gaussian data distribution), two-way analysis of variance (ANOVA) with Bonferroni's correction for multiple comparison and one-way ANOVA using Bonferroni's correction for multiple comparisons where applicable. Normality of data distribution was examined using Shapiro–Wilk's normality test. Statistical analysis was performed using GraphPad Prism (v9) software. Unclassified principal component analysis (PCA) was performed using a PCA toolbox in Matlab (*6*). Data was autoscaled prior to PCA calculation using venetian blinds cross validation and 5 cv groups (*7*). Classified discriminant analysis was performed using a classification toolbox in Matlab using linear discriminant analysis, bootstrap validation and 100 iterations. Hierarchical cluster analysis was performed in Matlab using the function clustergram. Differences were considered significant when $P < 0.05$. Differentially expressed genes were visualised with enhanced volcano plots implemented in R-Studio (*8*).

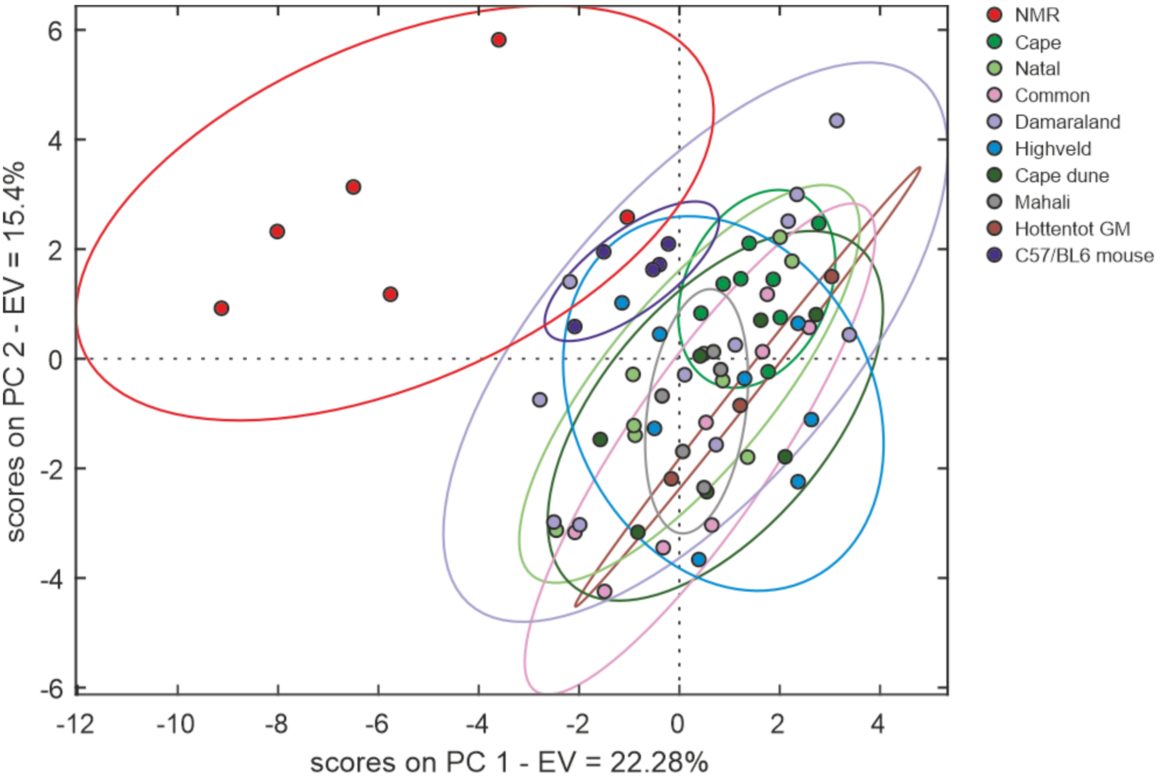


**Fig. S2.** Principal Component Analysis (PCA) of $^{1}H$ nuclear magnetic resonance metabolomic data showing separation of the NMR metabolomics data compared to all other genera. (n=3-11/group)

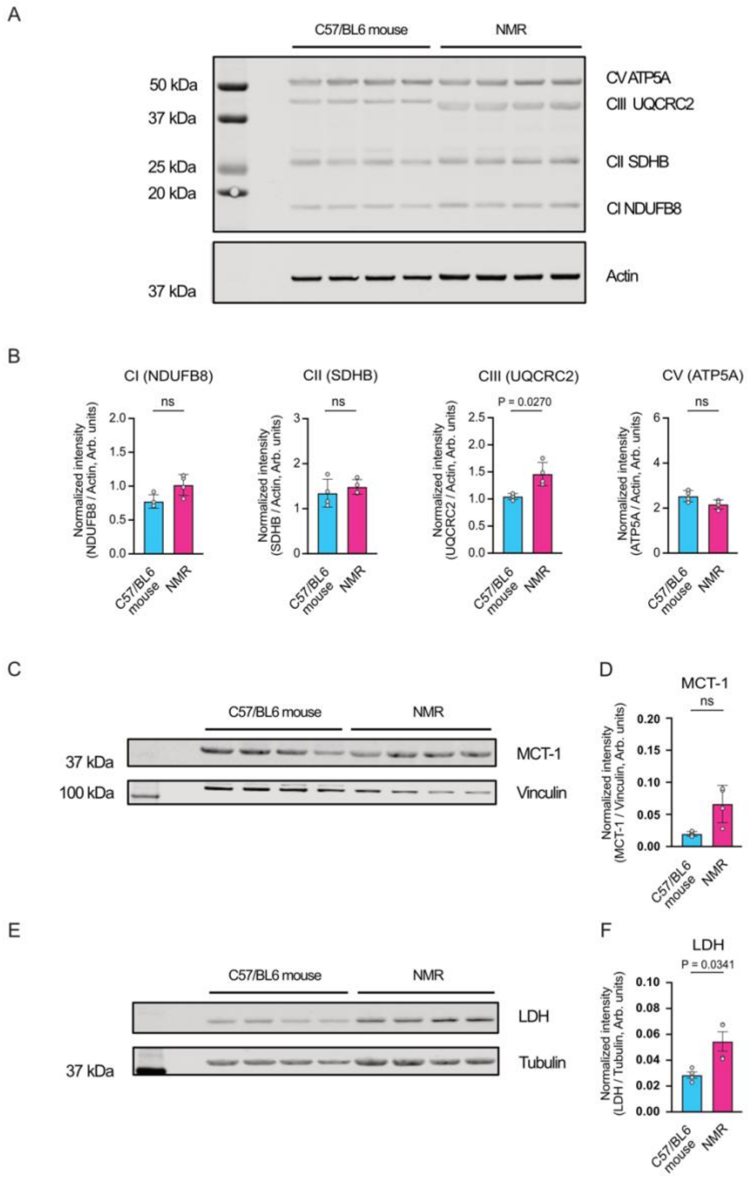


**Fig. S3.** Representative western blots (n = 4 animals/group) for the assessment of protein expression of mitochondrial electron transport chain (ETC) complexes, lactate dehydrogenase (LDH) and monocarboxylate transporter 1 (MCT-1) (**A**) Western blot of indicative subunits (ATP5A, UQCRC2, SDHB and NDUFB8) of mitochondrial ETC protein complexes (Complex I, II, III, and V) isolated from C57/BL6 mice and NMR hearts. (**B**) Quantification of subunits of oxidative phosphorylation complexes in C57/BL6 and NMR hearts. (**C**) Representative MCT-1 western blot from C57/BL6 mouse and NMR heart. (**D**) Quantified levels of MCT-1 protein from C57/BL6 mice and NMR hearts. (**E**) Representative LDH western blot from C57/BL6 mouse and NMR heart. (**F**) Quantified levels of LDH protein from C57/BL6 mice and NMR hearts. All experimental groups contain 4 animals per group and western blot analysis was performed using several technical replicates; protein expression normalised to loading control (details on y axis). Statistical analysis was performed using unpaired t-test (with Welch's correction).

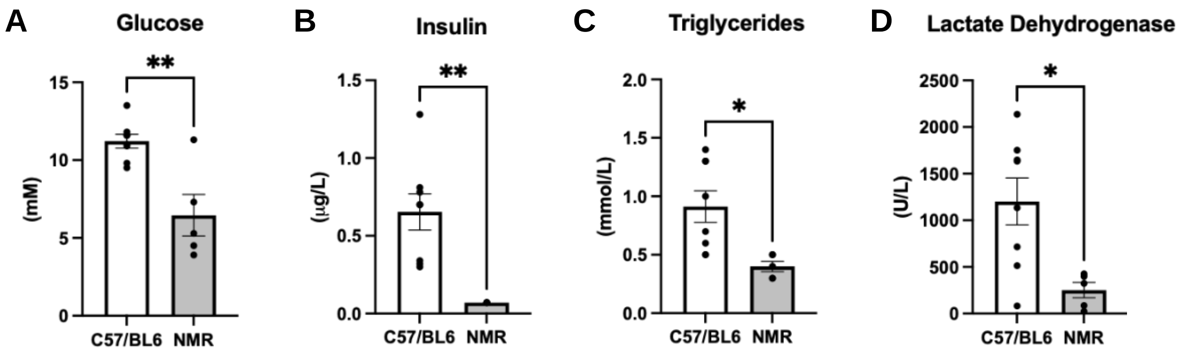


**Fig. S4.** Plasma metabolite concentrations of (**A**) glucose, (**B**) insulin, (**C**) triglycerides and (**D**) lactate dehydrogenase in NMR versus C57BL6 control mouse (NMR, n = 4 animals / group; C57BL6 (n = 8 animals / group), data mean + SEM, ** P<0.01 t-test.

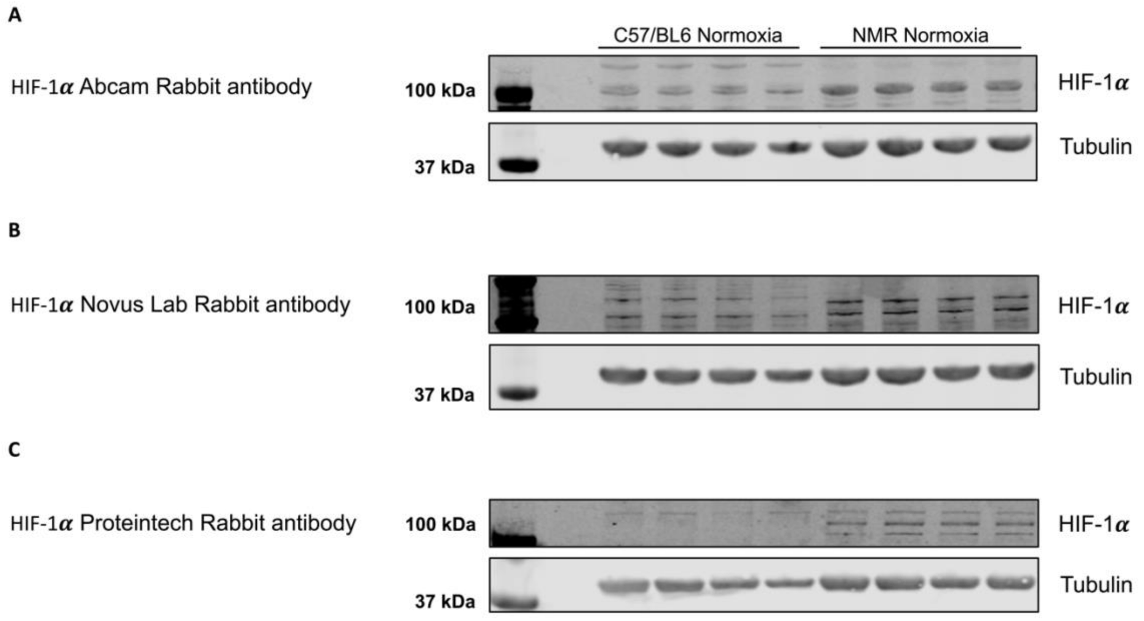


**Fig. S5.** Representative HIF-1α western blots using three different antibodies against different HIF-1α protein epitopes in cardiac tissue from C57/BL6 mouse and NMR (n = 4 animals / group). (**A**) Hif1α Abcam rabbit Ab, (**B**) Hif1α Novus lab rabbit Ab, (**C**) Hif1α Proteintech rabbit Ab.

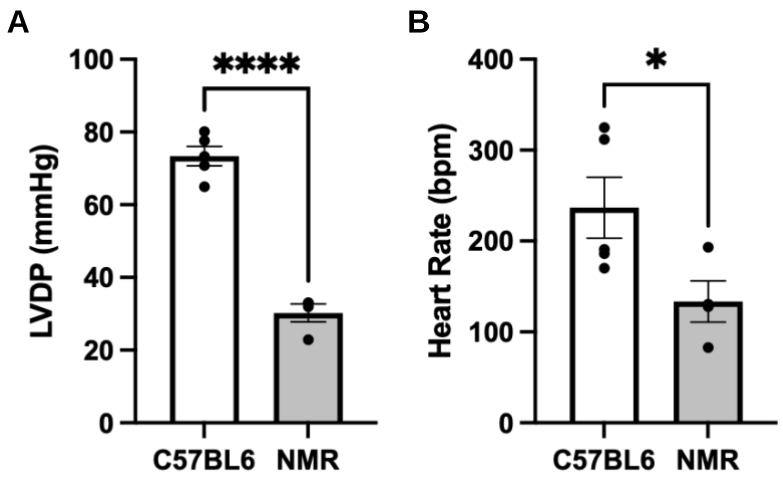


**Fig. S6.** *Ex vivo* cardiac function in Langedorff perfused hearts of NMR (n = 4 animals / group) and C57BL6 mouse (n = 5 animals / group), (**A**) left ventricular developed pressure (LVDP, mmHg) and (**B**) heart rate (HR, bpm), *** P<0.0001 vs control by t test.

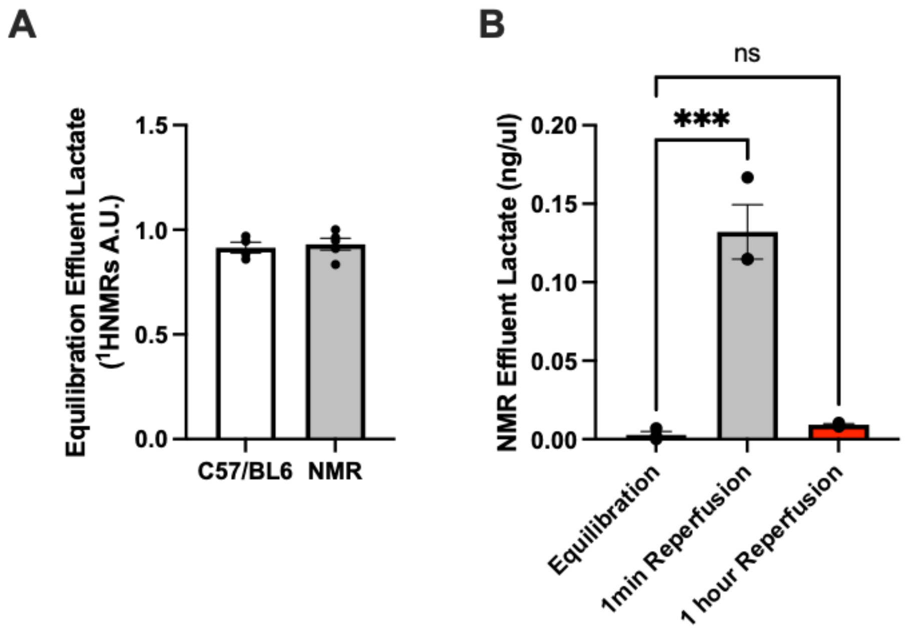


**Fig. S7.** Myocardial effluent lactate production (**A**) in NMR vs C57/BL6 mouse during equilibration (perfused heart coronary effluent collection) and (**B**) analysis of lactate concentration in coronary effluent of naked mole rat heart at equilibration, 1 minute and 1 hour after reperfusion. Two group comparison by t-test, multiple group comparison by one-way ANOVA. Data normality tested by Shapiro-Wilk. *P<0.05, **P<0.01 *** P<0.001 *** P<0.0001. (n = 3-5 animals / group)

| ***Species*** | ***Common name*** | ***Sample size*** | ***Heart Weight (mg)*** | ***Body weight (g)*** | ***Maximum lifespan (years)*** *(24)* | ***Age (years)*** |
|---|---|---|---|---|---|---|
| *Georychus capensis* | Cape mole rat | n=8 (2m, 6f) | 78.6±8.7 | 154±16.9 | 11 | 5 |
| *Bathyergus suillus* | Cape-dune mole rat | n=7 (5m, 2f) | 88.1±15.9 | 639±59.8 | >6 | 4-5 years |
| *Cryptomys hottentotus hottentotus* | Common mole rat | n=8 (6m, 2f) | 82.6±7.9 | 84.2±6.9 | 11 | 4-5years |
| *Cryptomys hottentotus pretoriae* | Highveld mole rat | n=8 (3m, 5f) | 81.9±10.4 | 102.7±11.1 | 11 | 4-5 years |
| *Cryptomys hottentotus mahali* | Mahali mole rat | n=5 (1m, 4f) | 85.9±21.8 | 114.8±8.6 | 11 | 4 years |
| *Cryptomys hottentotus natalenesis* | Natal mole rat | n=9 (3m, 6f) | 61.3±4.8 | 82.3±6.7 | 11 | 4 |
| *Fukomys damarensis* | Damaraland mole-rats | n=11 (3m, 8f) | 93.1±8.8 | 124.6±8.2 | 16 | 6-7 |
| *Heterocephalus glaber* | Naked mole rat | n=6 (4m, 2f) | 54.8±5.8 | 37.7±4.9 | >37 | 5 |
| *Amblysomus hottentotus* | Hottentot golden mole | n=3 (1m, 2f) | 76.3±9.9 | 58.6±8.6 | >1 | Adult (>1year) |
| *Mus musculus* | *C57/BL6 mouse* | n=5 (5m) | 155.2±0.1 | 25±1 | 2-3 | 0.1 |

**Table S1.** Morphological and life history characteristics of the African mole rat genera. m=male, f=female

**Data S1 file description**

Excel file containing a normalised comparison of the 30 most expressed genes in the C57/BL6 mouse and all the subterranean species examined in this study. Genes are ranked in descending order according to FPKM values (fragments per kilobase of transcript per million mapped reads). Mole-rat and golden mole gene descriptors are based on the nearest description identified. Yellow filled cells are unique to the naked mole-rat top 30. Common colour fills represent shared genes.

**Data S2 file description**

Excel file containing DEG data from RNA sequencing experiment (see supplementary methods above). Each sheet within workbook shows pairwise DEGs between NMRs and each other subterranean species in the data set, together with Log2 fold changes, P-values and gene details.
